# Fast prediction of protein flexibility

**DOI:** 10.64898/2025.11.30.691417

**Authors:** Jure Pražnikar

**Affiliations:** Faculty of Mathematics, Natural Sciences and Information Technologies, University of Primorska, Glagoljaška 8,6000, Koper, Slovenia; Department of Biochemistry, Molecular and Structural Biology, Institute Jožef Stefan, Jamova 39, 1000, Ljubljana, Slovenia

**Keywords:** protein flexibility, graphlets, predictive model

## Abstract

Advances in hardware have made molecular dynamics (MD) simulations of protein structures faster and more accessible to the scientific community. However, accurately estimating protein flexibility using MD remains computationally demanding, especially for large systems and long time scales. Several MD-based resources – including MdMD, the DynamD database, and more recently ATLAS and mdCATH – now provide MD trajectories for thousands of proteins, enabling the development of predictive models. Here, the Graphlet Degree Vector (GDV) is introduced as a lightweight, fast, and easy-to-implement linear model for predicting protein flexibility directly from atom coordinates. GDV is a 15-dimensional feature vector that captures local packing and the spatial connectivity of each atom with its nearby neighbors. Trained on a subset of globular-like proteins from the ATLAS database, the GDV model achieves a Spearman correlation of 0.828 compared to MD data. The model trained on ATLAS dataset was further evaluated on independent Nuclear Magnetic Resonance and cryo-electron microscopy datasets, demonstrating the robustness and generalizability of the GDV-based approach. A key advantage of the GDV model is that it requires no additional external or experimental data and can be applied in near real time (on the order of 10 seconds) even for large proteins with 20,000 atoms on a standard desktop or laptop. Overall, the results show that a lightweight, fast, and purely coordinate-based model can provide accurate and generalizable predictions of protein flexibility across diverse folds and sizes. The source code is available in the GitHub repository https://github.com/jure-praznikar/FastProtFlex. The data required for model training are available at https://doi.org/10.5281/zenodo.17771418.

## Introduction

Proteins are among the most abundant organic molecules in living systems and serve as fundamental building blocks of life. They perform nearly every essential cellular task, including catalyzing biochemical reactions, transmitting signals, and providing structural support, making them indispensable for proper organism function. Proteins are dynamic macromolecules, and their flexibility is closely linked to biological function. Therefore, studying protein flexibility is essential for understanding protein function. Molecular dynamics (MD) simulation is a fundamental and widely accepted computational method for characterizing protein motions at atomic resolution. However, MD is computationally demanding, especially for large systems or long timescales, which limits its routine use on standard desktop or laptop hardware. Despite hardware advances that have made MD simulations more accessible in the past decade, large-scale and real-time analysis of protein flexibility remains a challenge. To address these limitations, faster methods have been developed, including normal mode analysis Brooks et al. (1995); Case (1994); Ma (2005); Skjaerven et al. (2009) and various coarse-grained models, e.g. Gaussian Network Model Bahar et al. (1997); Haliloglu et al. (1997); Tirion (1996); Yang et al. (2009). These approaches trade some accuracy for computational efficiency and typically rely on physical approximations rather than data-driven learning.

Recently, the ATLAS database was introduced Vander Meersche et al. (2024), featuring standardized MD simulations, using CHARMM36m force field Huang et al. (2017), for over one thousand proteins of various sizes and folds. Because all simulations were generated using a unified protocol, ATLAS is free from user-specific choices and force field heterogeneity, making it especially suitable as a benchmark and training resource for predictive models of protein flexibility. From MD trajectories, atomic root-mean-square fluctuation (RMSF) profiles can be derived and used as target variables for model development.

In addition to MD databases, experimental databases such as Nuclear Magnetic Resonance (NMR) and cryo-electron microscopy (cryo-EM) are also important, as they provide information about protein flexibility. Currently, cryo-EM accounts for a significant portion of newly deposited macromolecular structures in the Protein Data Bank (PDB), and its experimental data are an important source of structural and flexibility information for macromolecules. Recently, cryo-EM datasets have been used to train deep learning models that predict protein flexibility directly from atomic coordinates and experimental electron density Song et al. (2024). This deep learning model demonstrates strong performance, and even very large proteins (25,000 atoms) can be processed in minutes.

In addition to experimental methods, artificial intelligence tools such as AlphaFold2 Jumper et al. (2021) have made remarkable progress and now enable highly accurate predictions of protein structures. These tools now generate high-quality 3D models directly from sequence data and provide per-residue confidence scores (pLDDT) to indicate prediction reliability. It has been shown that protein sequence determines not only the three-dimensional structure but also its flexibility Guo et al. (2022). Moreover, pLDDT scores show moderate to high correlations with MD- and NMR-derived RMSFs Vander Meersche et al. (2025). Furthermore, the pLDDT score has been successfully incorporated into software such as CABS-flex Wróblewski and Kmiecik (2024), where it improves flexibility prediction. Together, these developments suggest that protein flexibility can be effectively learned from structure-based features. Thus, the growing availability of MD protein trajectories, which is expected to continue to expand, offers a promising opportunity to develop models to predict protein flexibility Meyer et al. (2010); Van Der Kamp et al. (2010); Vander Meersche et al. (2024); Mirarchi et al. (2024).

Motivated by the availability of ATLAS data and the previous success of the Graphlet Degree Vector (GDV) linear model for B-factor prediction Pražnikar (2023, 2025) which was trained on a subset of the PDB-REDO database Joosten et al. (2009, 2014), this work extends the GDV linear model to predict protein flexibility at the atomic level. Graphlets– small, connected, induced, non-isomorphic subgraphs of a larger graph – are used to construct a feature vector for each atom, capturing the complexity of its local connectivity Pržulj (2007). In this approach, each protein structure is represented as a graph, with each node corresponding to an individual atom and edges containing information about the close contacts of each node. The atomic GDV is calculated from the larger graph and serves as a feature vector capturing information about local atom packing. In this study, the GDV linear model was trained on globular-like proteins from the ATLAS database. Cross-validation and evaluation on independent NMR (140 proteins) and cryo-EM (321 proteins) datasets demonstrated generalization and good predictive performance. The results presented in this work demonstrate that a lightweight, fast, and purely coordinate-based model can provide accurate predictions of protein flexibility across various folds and sizes.

## Methods

### Graphlet Degree Vector

Each protein structure was represented as a graph *G* = (*V, E*), where nodes *V* correspond to atoms and edges *E* connect pairs of atoms whose interatomic distance is less than a predefined cutoff of 7 Å. This approach convertsa three-dimensional atomic model into a network of atoms. The local packing of atoms is described using graphlet orbits. Graphlets are small, connected, non-isomorphic induced subgraphs of sizes 2, 3, and 4. Graphlets contain several automorphism orbits, which represent distinct topological roles of a node within the graphlet. There are 15 such orbits in total (labeled 0–14), capturing unique connectivity patterns (Figure 1A). For each atom, the number of times it participates in each orbit is counted, yielding a 15-dimensional GDV. The resulting *N ×* 15 matrix, where *N*is the number of atoms, describes the local network topology around each atom. Figure 1A shows all eight graphlets up to size four, while Figures 1B and 1C show an example of a 10-node graph and its corresponding GDV matrix.

**Figure 1.**
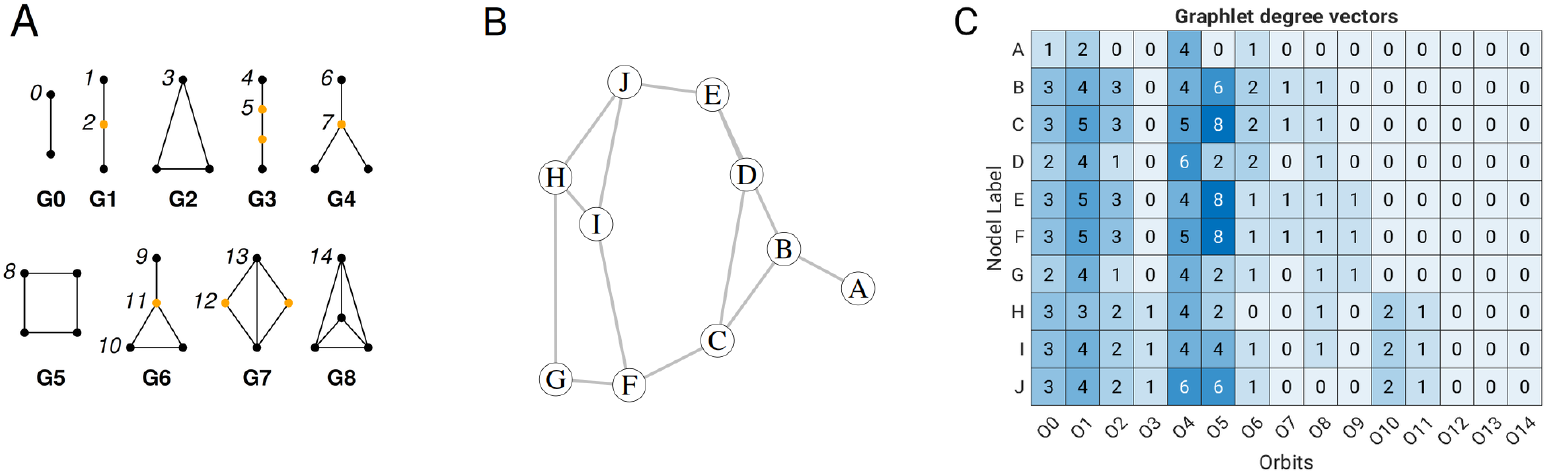
(A) Graphlets up to size 4 with corresponding orbits (0–14). (B) Example graph with 10 nodes and 13 edges. (C) Corresponding GDV matrix showing the orbit count for each node.

### GDV Linear model

A multiple linear regression model was used to predict atomic flexibility from GDV features. The dependent variable was the RMSF of atoms, and the explanatory variables were the GDV orbit counts. The multiple linear model is written as follows:

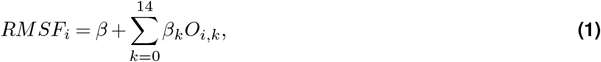

where *RMSF*_*i*_ is the observed fluctuation of atom *i*; *β* is the intercept; *β*_*k*_ are the regression coefficients for orbit *k*; and *O*_*i*,*k*_ are the GDV components for atom *i*. Both the dependent and explanatory variables were log-transformed and normalized (zero mean, unit standard deviation) within each protein. Model parameters were estimated using ordinary least squares regression. Predictive performance was assessed using 10-fold cross-validation by computing the Spearman correlation between MD–derived *RMSF* values and those predicted by the GDV linear model. The correlation was calculated at the residue level, specifically for C*α* atoms. This procedure was repeated across three MD replicates presented in the ATLAS database to assess robustness and generalization. The resulting averaged coefficients *β*_*k*_ define the final GDV-based linear model.

## Data

### ATLAS data set - globular-like proteins

The ATLAS database (November 2024 release) contains 1,938 non-redundant protein chains. For each protein, standardized MD simulations were performed using three different random seeds to capture a broad range of conformational motions. For GDV model development, a subset of ATLAS entries was selected based on the following criteria: (i) the protein belongs to CATH classes 1–3 Orengo et al. (1997); (ii) the number of residues exceeds 50; (iii) the radius of gyration is less than *N* ^0.6^ (to ensure a globular-like structure); and (iv) the TM-score Zhang and Skolnick (2005) between the initial and final conformations, as well as between the most divergent conformers, exceeds 0.7. The TM-score threshold ensured inclusion of proteins that do not undergo substantial fold changes during MD simulations. Applying these filters yielded a subset of 1,052 ATLAS entries. For additional benchmarking, the first release of the ATLAS dataset (November 2023), which contains 1,390 protein chains, was also used.

### NMR dataset

The NMR protein benchmark set was used and consists of 140 non-redundant protein structures Jamroz et al. (2012). Because NMR ensembles consist of multiple conformers, a critical consideration is which structure to use for RMSF prediction. The conformer most similar to the average NMR structure, defined as the model with the minimum C*α* r.m.s.d. relative to the mean coordinate structure, was identified and used as the representative three-dimensional model for prediction.

### Cryo-EM dataset

In addition to the ATLAS and NMR dataset, the cryo-EM benchmark data set Song et al. (2024) was also used. This high-quality, non-redundant protein dataset comprises 335 entries compiled for training and evaluating RMSF-net, focusing on cryo-EM maps with 2–4 Å resolution. All proteins were subjected to molecular dynamics simulations using the AMBER force field Case et al. (2023, 2005). The dataset was obtained from the open-access Figshare repository https://doi.org/10.6084/m9.figshare.25893670, (Nov. 2025). Upon inspection, 14 models contained NaN values in the B-factor column of the PDB files and were excluded, resulting in 321 proteins used for analysis.

### Software and Hardware

The R package, version 4.5.1 R Core Team (2025), was used for data analysis with the following packages: orca, version 1.1-3 Hočevar and Demšar (2014, 2016); netdist, version 0.4.9100 O’Reilly (2024); bio3d, version 2.4-5 Grant et al. (2006); igraph, version 2.2.0 Csardi and Nepusz (2006); caret, version 7.0-1 Kuhn (2008); pdist, version 1.2.1 Wong (2022). R scripts are available at: https://github.com/jure-praznikar/FastProtFlex. The experiments were conducted on a system with a 13th Intel Core i9-13900 processor, 5200 MHz RAM, and the openSUSE 15.6 operating system.

## Results

### The performance of linear GDV model

A linear GDV model was trained and validated on three independent ATLAS globular-like protein datasets (replicates 1–3). For each replicate, 10-fold cross-validation was performed, and model performance was evaluated using the Spearman correlation between MD-derived and predicted RMSF values. Figure 2 shows the 15 regression coefficients obtained for each dataset, demonstrating their consistency across replicates. The averaged coefficients (eq. 2) define the final GDV model, representing the consensus linear relationship between GDV features and RMSF across all three replicates:

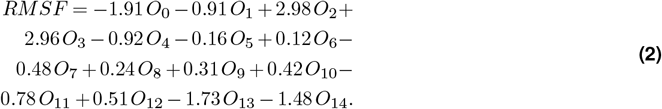

**Figure 2.**
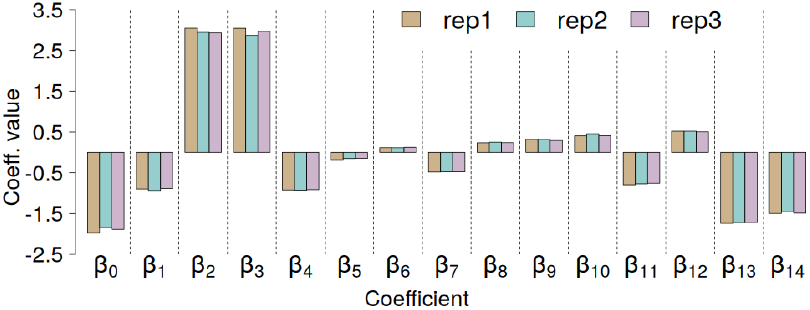
GDV linear model regression coefficients for three independent replicates from the ATLAS data set.

The performance of the GDV linear model trained on the ATLAS globular-like dataset (N = 1,052) using three independent replicates is shown in Figure 3A. The mean Spearman correlations between the predicted and MD-derived RMSF values were 0.794, 0.793, and 0.792 for replicates 1, 2, and 3, respectively (Figure 3A). When the best-performing replicate was selected for each protein, the mean correlation increased to 0.828.

**Figure 3.**
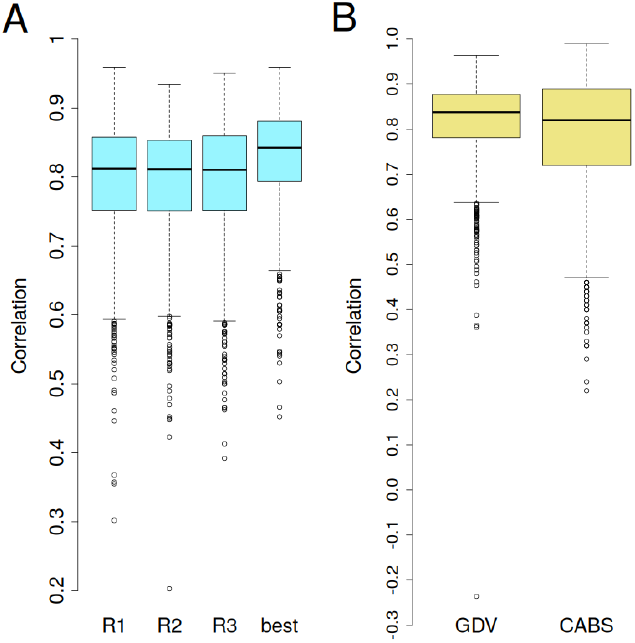
(A) Boxplots of correlations between MD-derived and GDV-predicted RMSF for each of the three replicates and for the best-performing replicate per protein for the globular-like ATLAS subset. (B) Boxplots of the best-selected correlations (among three replicates) for the GDV model and CABS-flex on the full ATLAS dataset (November 2023 release).

### Benchmark Part 1 – ATLAS Dataset

To benchmark the GDV linear model, the full ATLAS database (November 2023 release) was used and compared with CABS-flex, which integrates AlphaFold pLDDT scores Wróblewski and Kmiecik (2024). In that study, the largest protein (PDB ID: 6SUP), containing approximately 17,000 heavy atoms, was excluded. CABS-flex is a coarse-grained simulation tool that models protein flexibility using knowledge-based statistical potentials and Monte Carlo dynamics Jamroz et al. (2013). In its default restraint mode, it achieved a correlation of 0.661; however, when the pLDDT score was appropriately incorporated as an additional feature, the correlation increased to 0.793 Wróblewski and Kmiecik (2024). The GDV model achieved an average correlation of 0.817 (Figure 3B), which is slightly higher than the average correlation obtained with CABS-flex. Notably, the GDV linear model is based exclusively on atomic coordinates and does not incorporate additional features such as secondary structure or AlphaFold2-derived pLDDT scores, in contrast to CABS-flex.Beyond accuracy, a key motivation of this work was to develop a fast, lightweight algorithm suitable for personal computers and laptops. Therefore, CPU time and RAM usage during RMSF prediction using the GDV linear model were evaluated. The most computationally demanding step is graph construction, which requires calculating all pairwise atomic distances. This operation also drives memory consumption, as it involves generating a large two-dimensional distance matrix.Figure 4A shows the relationship between protein size and peak RAM usage. The dependency is non-linear, approximately quadratic. For proteins with about 8,000 atoms, RAM usage reached 4 GB. Proteins with up to 4,000 atoms were converted to graphs in less than one second each (Figure 4B). For larger systems, however, converting a 3D model to a graph required only about three seconds, even for proteins with 8,000 atoms. Extrapolating the RAM-to-protein size relationship shown in Figure 4A suggests that for very large proteins (approximately 40,000 atoms), RAM rather than CPU time becomes the limiting factor on standard desktop or laptop computers and requires more than 100 GB of memory. To address this, a by-part GDV calculation scheme was implemented and compared to the standard all-at-once approach. The protein of interest is divided into smaller, non-overlapping segments (e.g., three parts of 100, 100, and 98 residues for a 398-residue protein). For each segment, surrounding residues within 12 Å of C*α* atoms were included, typically forming subsets of about 2,200 atoms (approximately 750 core and 1,500 neighboring atoms). GDVs were then computed for each segment and concatenated to yield the complete matrix. Figure 4C,D illustrates RAM usage and CPU time with the by-part approach, which was applied only to proteins exceeding 2,000 atoms. This optimization reduced the memory footprint to below 1 GB and yielded a nearly linear relationship between computation time and protein size. These results demonstrate that using the by-parts approach to compute the GDV matrix keeps RAM usage low and CPU time relatively short, allowing the GDV model to run on standard laptop computers even for large proteins in near real time.

**Figure 4.**
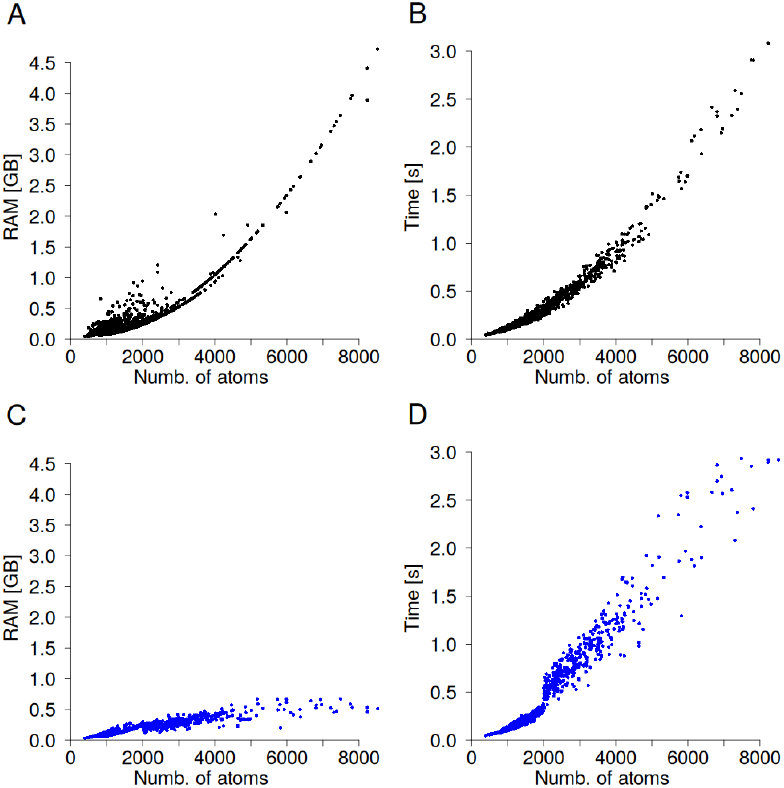
(A, C) Number of atoms versus peak RAM usage, and (B, D) number of atoms versus computation time for two calculation strategies. Panels (A) and (B) correspond to the all-at-once approach, while panels (C) and (D) show results for the by-parts approach.

### Benchmark Part 2 - Comparison to NMR ensembles

To assess how RMSF values derived from NMR ensembles align with those predicted by the GDV model, multiple NMR conformers of a selected protein (PDBid: 1S6L) were analyzed. Because each conformer has distinct atomic coordinates, individual RMSF predictions differ slightly across conformers. Figure 5A presents NMR-derived alongside predicted RMSF values for 20 conformers. Strong agreement was observed, with Spearman correlation coefficients ranging from 0.80 to 0.89 across the twenty conformers. To extend the analysis, correlations werealso calculated for the NMR benchmark dataset comprising 140 proteins. Figure 5B presents the distribution of correlations, with a mean (median) value of 0.729 (0.776), comparable to the accuracy reported for CABS-flex Jamroz et al. (2014). These results further demonstrate the robustness of the GDV model, which, despite being trained exclusively on ATLAS dataset, shows strong agreement with fluctuations observed in NMR ensembles.

**Figure 5.**
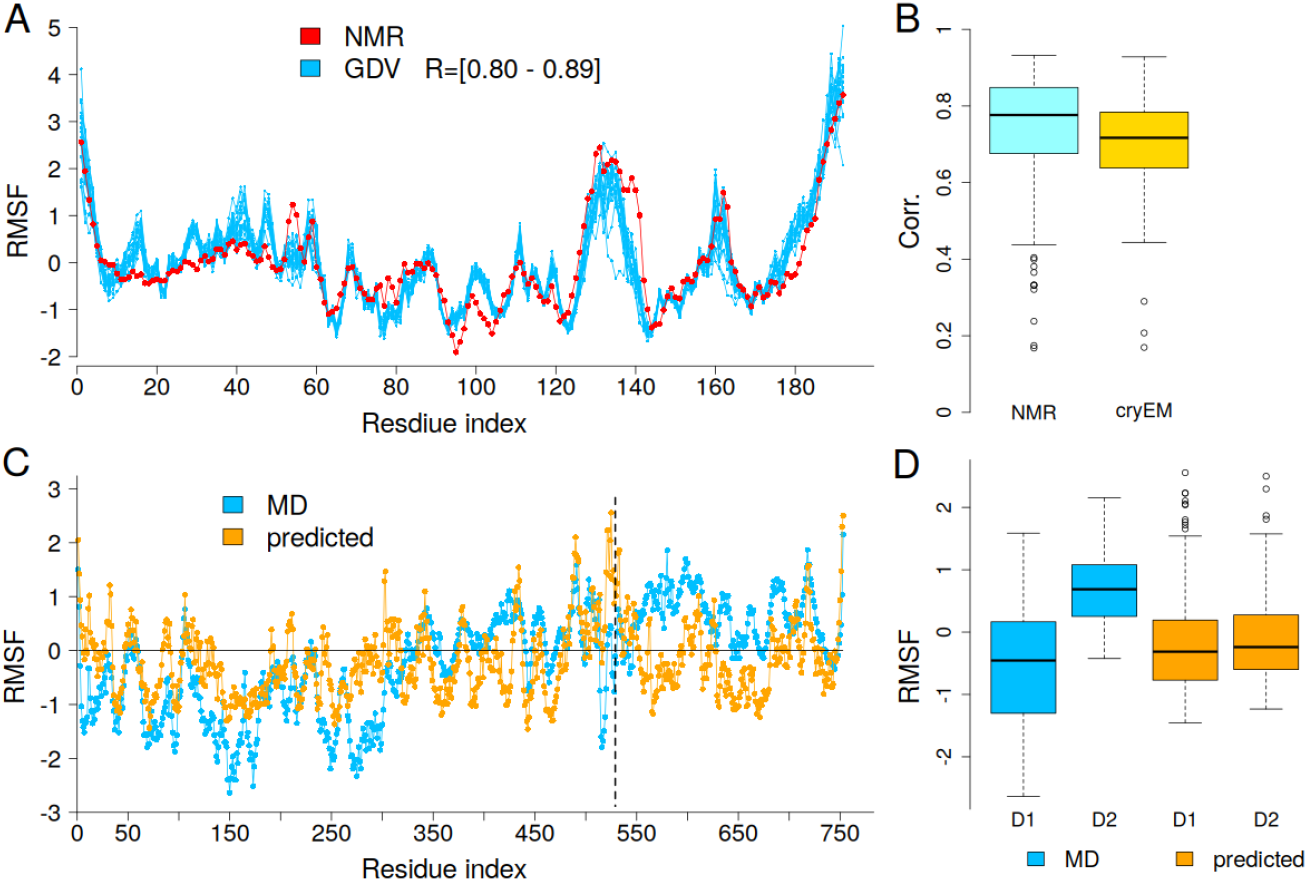
(A) GDV-predicted RMSF profiles for 20 NMR conformers and NMR-derived RMSF for PDBid: 1S6L. (B) Boxplots of correlations between GDV-predicted RMSF and NMR- and cryo-EM–derived RMSF values. (C) MD-derived and GDV-predicted RMSF profiles for PDBid: 1C96. (D) Boxplots showing RMSF distributions for the two domains: domain 1 (D1, residues 1–528) and domain 2 (D2, residues 529–753), corresponding to SCOP domains c.83.1.1 and c.8.2.1, respectively.

### Benchmark Part 3 - Cryo-electron microscopy

The cryo-EM dataset includes 335 proteins Song et al. (2024), for which MD simulations were performed using the AMBER force field Case et al. (2023, 2005). This dataset contains significantly larger proteins, with an average size of ~1,700 residues, compared to the ATLAS dataset, which has a mean size of approximately *~ 2*50 residues. The mean correlation between MD and predicted RMSF was 0.704 (Figure 5B), which is lower than the 0.765 and 0.742 achieved by the RMSF-net and RMSF-net_pdb neural network models Song et al. (2024), respectively. However, it is important to note that GDV linear model does not incorporate experimental data, such as electron density maps, unlike RMSF-net. However, these results demonstrate the generalizability of the GDV linear model, which was trained on the ATLAS dataset and still provided satisfactory predictions for independent cryo-EM dataset.

### Study Cases

#### Non-globular protein

For the GDV model, one clear outlier shows a markedly low correlation of −0.236 (see Figure 3B). This protein corresponds to PDBid: 4KE2 (chain C), which was excluded from the globular-like subset due to its large radius of gyration relative to its size of 192 residues Hong and Lei (2009); Tanner (2016); Pražnikar (2021), indicating a non-globular topology. Indeed, this is an alanine-rich antifreeze protein that forms a rod-like dimeric structure composed *α*-helices arranged into a four-helix bundle Sun et al. (2014). Thus, for this protein, the results indicate that the GDV linear model does not accurately capture fluctuations caused by *α*-helix bending.

#### The inter-domain domain flexibility

PDB entry 1C96 (chain A) shows relatively low correlations between predicted and MD-derived RMSF values; 0.46, 0.50, and 0.58 for replicates 1-3, respectively. This protein comprises two domains, classified in Structural Classification of Proteins (SCOP) Andreeva et al. (2013) as c.83.1.1 (residues 1–528) and c.8.2.1 (residues 529–753). The GDV linear model tends to overestimate fluctuations in the larger domain while underestimating them in the smaller one (Figure 5C). Boxplots in Figure 5D illustrate that MD simulations capture a pronounced difference in flexibility between domains, whereas this contrast is notably reduced in the GDVlinear model. This case highlights a limitation of the approach—its reduced sensitivity to large-scale, inter-domain motions.

#### Wrong model - poor prediction

As part of the preprocessing for MD simulations in the ATLAS database, missing residues (*≤* 5) were modeled using MODELLER Webb and Sali (2016). For example, in PDB entry 1BY2 (chain A), four C-terminal residues (Arg114–His117) were absent and modeled to connect Thr118 and Leu119 according to ATLAS residue numbering. Visual inspection and WHAT_CHECK Hooft et al. (1996) analysis revealed seven steric clashes (bumps), defined as atomic contacts closer than the sum of van der Waals radii minus 0.4 Å. These involved residue pairs Ser45–His117, Arg11–Trp31, Cys48–His117, Asp40–Leu41, Val46–Ser85, Cys48–Gly52 and Cys48–Thr118.

Figure 6A shows the initial model, with the modeled C-terminal tail highlighted in magenta, and snapshots at 25 ns, 50 ns, and 100 ns of the MD trajectory. During MD simulation, the tail (Ser115–Leu119) moves out of the cavity where it was initially placed. Both N- and C-terminal regions exhibit high flexibility in MD simulations (Figure 6B). In contrast, the GDV linear model underestimates fluctuations in the C-terminal tail and in the *α*-helical region spanning Asp40–Ala50. This discrepancy arises from the artificially high local contact density caused by the mispositioned C-terminal tail of initial model. It should be noted that the GDV model relies solely on atomic coordinates; consequently, any structural inaccuracies in the input model directly affect RMSF prediction quality. For the initial model, the correlation between MD-derived and predicted RMSF was 0.45, which increased to 0.72 when the final 100 ns snapshot was used (Figure 6B). A partially incorrect 3D protein model degrades both GDV-based predictions and the MD simulation, making the resulting MD-derived RMSF values less reliable.

**Figure 6.**
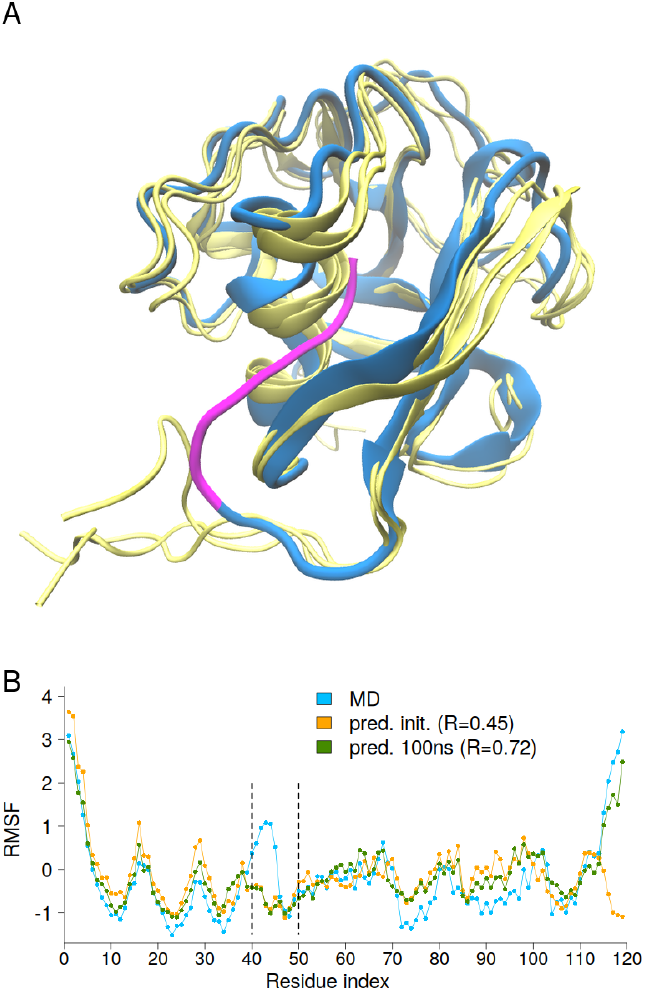
(A) Initial structure (blue) with the modeled C-terminal tail (Ser115–Leu119) highlighted in magenta, and MD snapshots at 25 ns, 50 ns, and 100 ns shown in yellow. (B) MD-derived and predicted RMSF profiles for PDBid: 1BY2, using coordinates from the initial model and the 100 ns snapshot.

#### Large protein system

PDB entry 6SUP was excluded from globular-like dataset because its radius of gyration exceeded value *N* ^0.6^, where *N* denotes the number of amino acids, indicating a non-globular structure. Interestingly, this protein was also omitted from the CABS-flex study Wróblewski and Kmiecik (2024) for a different reason—its large size (~17,000 heavy atoms), which makes computation substantially more demanding. Therefore 6SUP was used as a benchmark to assess both the speed and accuracy of the GDV linear model for large proteins. RMSF values were predicted using two approaches: (i) a single all-at-once calculation and (ii) a by-parts approach. Notably, protein 6SUP was divided into 22 *chunks*, each comprising a core region of ~100 residues and a surrounding region defined by cutoff distance (12-15 Å). All-at-once calculation required 12 s and ~19 GB of RAM, whereas theby-parts computation (cutoff 12 Å) was nearly twice as fast (6 s) and used less than 1 GB of memory (Table 1). The correlations with MD-derived RMSF were ~0.79 for C*α* atoms. Only minor differences were observed between the all-at-once and by-parts approaches (Table 1).

**Table 1.** Performance statistics for PDB entry 6SUP from the ATLAS database include correlation, computation time, and peak RAM usage.

|  | all-at-once | by-parts<br>12 Å | by-parts<br>15 Å |
| --- | --- | --- | --- |
| rep. 1 | 0.804 | 0.793 | 0.803 |
| rep. 2 | 0.790 | 0.786 | 0.791 |
| rep. 3 | 0.798 | 0.792 | 0.799 |
| time [s] | 12 | 6 | 8.5 |
| mem. [GB] | 19 | 0.7 | 1.1 |

As the cutoff distance defining the surrounding region of each segment increases, both accuracy and computation time increase; conversely, reducing the cutoff decreases both accuracy and computation time. This trend is shown in Table 2, which summarizes the results for the entire cryo-EM dataset using by-parts approach. Reducing the cutoff below 12 Å results in a notable drop in correlation, while values of 15 and 20 Å yield similar performance. Thus, a cutoff of 12–15 Å provides an effective balance between accuracy and computational efficiency. For the full cryo-EM dataset (335 proteins), the total computation time was several tens of minutes (Table 2). These results demonstrate that the GDV linear model provides accurate, near–real-time RMSF predictions even for very large proteins.

**Table 2.** Performance statistics (correlation and computation time) for the cryo-EM dataset. Graph conversion was performed on the full dataset of 335 proteins, while correlations were calculated for a subset of 321 proteins (see Methods for details).

| cutoff Å | mean corr. | time (minutes) |
| --- | --- | --- |
| 8 | 0.593 | 15 |
| 12 | 0.690 | 26 |
| 15 | 0.704 | 33 |
| 20 | 0.706 | 57 |

## Conclusion

In this study, the GDV linear model is introduced as a fast, lightweight and accurate method for predicting protein flexibility. A key advantage of this approach, beyond its simplicity and speed – which allow near real-time use even on standard laptops – is that it relies solely on atomic coordinates and does not require any additional external data or annotation library. Even for proteins with approximately 20,000 atoms, it takes less than 10 seconds to convert the 3D structure to a graph, calculate GDVs, and predict all-atom RMSF profile.

For any protein structure, the algorithm computes atomic GDVs, applies the GDV linear model, and predicts normalized RMSF values using only atomic coordinates. This allows efficient, interpretable, and reproducible estimation of atomic fluctuations without requiring additional input features such as atom type, secondary structure, or AlphaFold2-derived confidence scores. However, there are several opportunities to improve the GDV linear model. Performance could be enhanced for non-globular, rod-like proteins, and special attention should be given to inter-domain fluctuations. Currently, the model captures the local environment of each atom through 15 local features, which do not account for long-range conformational effects important for inter-domain motions.

The GDV linear model has previously been used to predict B-factors in crystal structures, including assessing the impact of large ligands such as DNA (deoxyribonucleic acid) and crystal packing. Similarly, GDV linear model could be applied to study RMSF profiles in monomeric versus homo-dimeric states, or in protein–protein and protein–DNA complexes, demonstrating its versatility for a range of structural biology applications.

## Author contributions statement

J.P. conceived of the presented idea, analysed the data, performed the calculations and wrote the manuscript.

## Acknowledgments

The author thank the anonymous reviewers for their valuable suggestions.

